# Fast & Fair peer review: a pilot study demonstrating feasibility of rapid, high-quality peer review in a biology journal

**DOI:** 10.1101/2025.03.18.644032

**Authors:** Daniel A. Gorelick, Alejandra Clark

## Abstract

Traditional peer review is slow, often delayed by the time-consuming process of identifying reviewers and lengthy review turnaround times. This study tests the feasibility of the Fast & Fair peer review initiative by evaluating whether we could implement and adhere to a structured timeline for rapid peer review at Biology Open. A 6-month pilot conducted from July to December 2024 evaluated a structured workflow with pre-contracted reviewers under two payment models: a freelance model or a retainer model. All manuscripts assigned to two of the journal’s ten academic editors were included in the Fast & Fair peer review initiative experiment. A structured editorial time-line ensured that all manuscripts received reviews and first decisions within 7 business days. The results demonstrated that 100% of Fast & Fair peer review initiative manuscripts met the turnaround target, with a mean of 4.6 business days (n=20 manuscripts). Review quality was maintained, as indicated by assessments by academic editors. The freelancer model outperformed retainers in cost-effectiveness. These findings suggest that the Fast & Fair peer review initiative is feasible and does not compromise review quality. While scalability remains to be tested, the initiative eliminates a major bottleneck in traditional peer review by streamlining reviewer identification and enforcing a strict editorial timeline.

## INTRODUCTION

Traditional peer review is a slow process, often taking weeks to identify reviewers and months before a first decision is reached (Nguyen et al., 2015). Various strategies have been explored to accelerate this process, including deadline adjustments, monetary incentives, and public recognition, with mixed results across disciplines.

One approach has been to impose shorter review deadlines. A study of manuscripts submitted to the Journal of Public Economics found that reducing the review deadline from 45 to 28 days led to a median decrease of 12 days in review completion time, suggesting that time constraints can serve as an effective nudge for reviewers to prioritise their assignments (Chetty et al., 2014). However, it remains unclear whether similar deadline enforcement would be as effective in biomedical sciences.

Monetary incentives have also been tested as a means of expediting peer review. In a 15-month study of manuscripts submitted to the Journal of Public Economics, offering reviewers a $100 reward for submitting a review within four weeks further reduced median review times by 8 days compared to deadline enforcement alone (Chetty et al., 2014) ($100 in 2010, the time of this study, is equivalent to the purchasing power of $146 today, https://www.bls.gov/data/inflation_calculator.htm).

Cotton et al. conducted a quasi-randomised trial at the medical journal Critical Care Medicine, finding that monetary incentives modestly increased the rate of peer review completion and reduced turn-around times by approximately 1 day without compromising review quality (Cotton et al., 2025). However, their incentive approach did not explicitly link payments to clearly defined performance expectations, potentially limiting the observed improvements.

The British Medical Journal (BMJ) offers all reviewers a free 12-month online subscription to one of the BMJ journals. Beginning in Jan 2025, BMJ offers patients and members of the public £50 for completing a review for The BMJ (Doble et al., 2024). However, it remains to be seen whether this compensation scheme will result in improved peer review.

In contrast, the concept of “motivation crowding” suggests that introducing monetary rewards may harm peer review. Motivation crowding refers to a psychological phenomenon in which external incentives, such as monetary rewards or public recognition, undermine intrinsic motivation for a task. Instead of enhancing performance, these incentives can lead to lower engagement or effort over time (Frey and Jegen, 2001). In the context of peer review, introducing monetary incentives may cause reviewers to perceive the task as a transactional obligation rather than an intrinsic academic duty, leading to lower review quality or reduced long-term participation.

Another incentive to improve peer review is public recognition for reviewers as offered by <u>ORCID</u>, <u>Reviewer Credits</u>, and <u>F1000Research</u>. Some journals have experimented with publicly posting reviewer turnaround times, leading to a modest reduction in review completion time, primarily among senior researchers (Yu et al., 2024). Platforms like <u>Publons</u> (now Web of Science Reviewer Recognition Services from Clarivate), introduced accolade awards for frequent reviewers (Preston and Johnston, 2013; Ravindran, 2016), but a large-scale study found that receiving such an award led to reviewers completing fewer reviews in subsequent years, possibly because they perceived their obligation as fulfilled or because the marginal value of additional accolades diminished over time (Yu et al., 2024). The negative impact was most pronounced among reviewers with high social capital and in developing countries, where academic recognition may carry different incentives.

Here, we sought to directly test the hypothesis that providing a financial incentive to reviewers could decrease turnaround time while still maintaining the quality of peer review in a biology/biomedical journal. We designed and implemented a system capable of rapidly reviewing manuscripts while maintaining high-quality reviews through editorial assessment. The goal of this pilot study was to evaluate whether such a system could be effectively implemented, assess adherence to a structured peer review timeline and determine the feasibility of rapid peer review. Additionally, the study examined the cost-effectiveness of different reviewer models and measured review quality through editorial scoring.

## MATERIALS AND METHODS

The Fast & Fair peer review initiative process followed a structured editorial timeline, where each manuscript was reviewed by two reviewers (Fig. 1). On the first business day after submission, all manuscripts underwent the standard evaluation before being assigned to an academic editor. Biology Open staff evaluated the manuscript for compliance with the rubric for publication (https://journals.biologists.com/bio/pages/rubric), including scientific soundness and potential indications of paper mill submissions. Biology Open staff also assessed eligibility for the trial: submissions in the subject areas handled by one of the two editors, and submissions that did not include peer review reports from other journals. Manuscripts that were transferred from other journals with reviews, including those from Review Commons, were excluded from the trial. If the submission was deemed valid, it was assigned to one of the two academic editors with relevant subject-area expertise.

**Fig 1.**
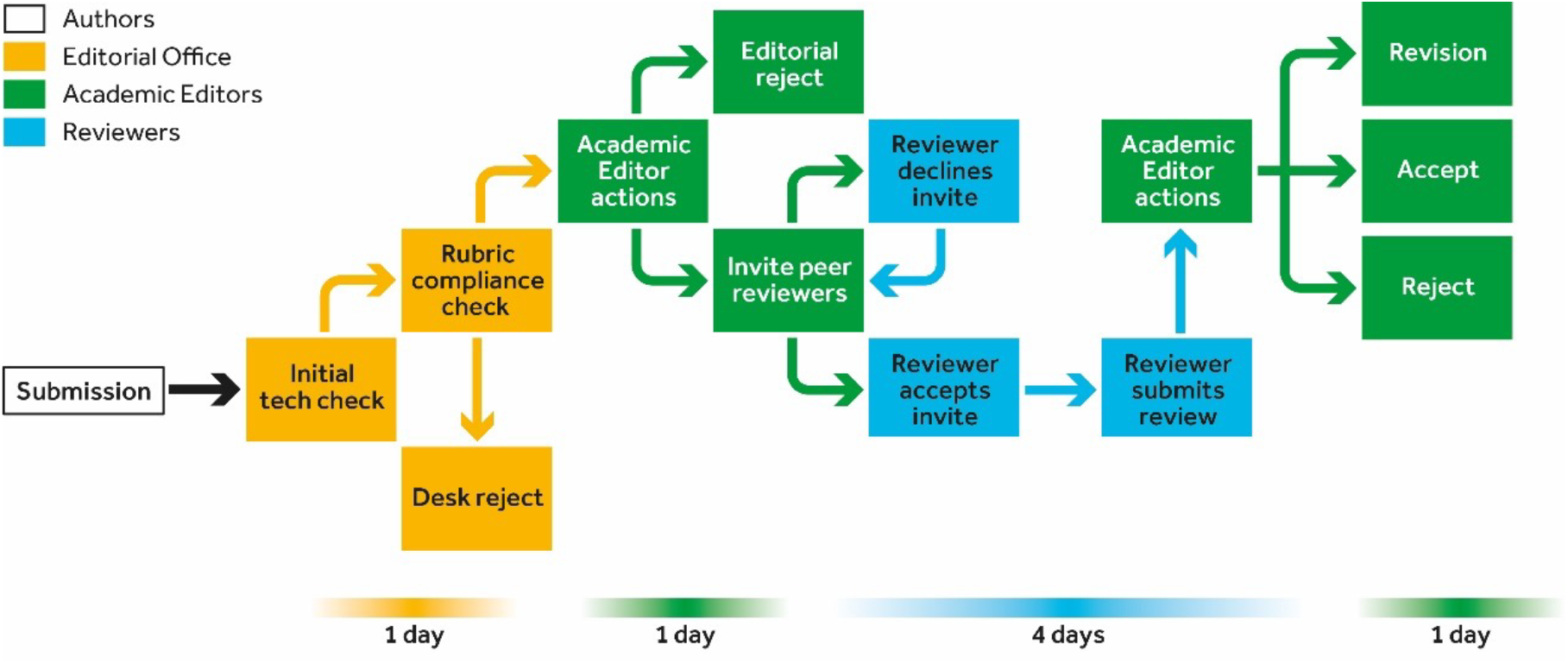
Editorial workflow for Fast & Fair peer review manuscripts. Initial checks (including rubric compliance) by editorial staff take place within 1 business day (yellow). Academic editors take an additional business day to decide if manuscripts are sent for peer review or desk rejected (green). If sent for peer review, reviewers are expected to reply to invites within 1 business day and complete their reviews within 4 business days of accepting (blue). Academic editors then take 1 business day to issue the first decision (revision, accept, reject).

By the second business day, the handling editor invited two peer reviewers. Freelancer reviewers were required to accept or decline the invitation within 1 business day, whereas retainer reviewers were obligated to accept all assigned review requests (see below, Reviewer models tested). Upon acceptance, reviewers were given 4 business days to complete their evaluations. On the final day of the process, the handling editor assessed the two submitted review reports and made a decision on the manuscript: accept, reject, or return to the authors for revisions. Authors were not notified of the Fast & Fair initiative and were kept blind as to whether their manuscripts were sent through conventional peer review or Fast & Fair.

### Pre-identifying reviewers

One of the primary challenges in traditional peer review is the difficulty in securing reviewers, which often leads to substantial delays. To overcome this hurdle, we pre-recruited reviewers who committed to a rapid response. Areas of reviewer expertise were chosen based on past submission trends. As of December 31, 2024, 80 reviewers were recruited through targeted outreach efforts, including invitations to previous Biology Open reviewers, invitations to <u>preLighters</u> (scientists who review and highlight preprints on a community platform supported by The Company of Biologists), and contacts within relevant research fields. These pre-recruited reviewers were then categorised into either the freelancer or retainer model to support timely peer review.

### Reviewer models tested

Two reviewer payment models were tested during the pilot. Retainer reviewers received £600 per quarter in return for reviewing three manuscripts per quarter. They were required to accept all assigned review requests and were paid £600 regardless of whether they reviewed zero, one, two or three manuscripts per quarter.

Freelance reviewers, in contrast, were paid £220 per manuscript. Freelancers had one business day to accept or decline review invitations (for any reason), but, if the invitation to review was accepted, freelancers were required to submit their reports within 4 business days.

Reviewer expectations were outlined in a contract. Both sets of contracts contained the same terms and conditions with regards to quality standards and compliance with journal reviewer guidelines. Both sets of contracts stipulated that if time or quality standards were not met, reviewers would not be paid.

### Review quality assessment

To ensure that review quality was not compromised, academic editors rated each review on a discrete three-level scale. A score of 100 indicates that the review is helpful and adequate, a score of 50 indicates that the review is helpful but needs improvement, a score of 0 indicates the review is unhelpful. If a reviewer were to receive a score of 0 they would not be paid nor be invited to review again under the initiative, whereas those scoring 50 would receive feedback and guidance for improvement. In addition, to ensure that the fast turnaround times of the Fast & Fair editorial workflow did not compromise the quality of editorial assessment, all manuscripts received internal oversight from editorial staff.

## RESULTS

The Fast & Fair peer review initiative pilot was implemented for all manuscripts handled by two of Biology Open’s ten academic editors: one editor, who handles manuscripts related to mammalian development, stem cells, growth regulation, mitochondrial biology, cell metabolism, cell signaling and cell death; and another editor, who handles manuscripts in the fields of accelerometry, anthropology, behavior, comparative physiology, diving, ecophysiology, energetics and metabolism. All other manuscripts continued under conventional peer review and therefore acted as a control for the pilot.

The journal received 25 eligible manuscripts for the trial (i.e. in subject matters within the expertise of the two academic editors). Three manuscripts were editorially rejected without being sent for peer review. 20 manuscripts were sent for peer review (mean time from submission to decision, with reviews, 4.6 days, range from 2 to 7 business days, Fig. 2). Two manuscripts were excluded from the experiment due to lack of scientific expertise in the pool of contracted reviewers and were handled via conventional peer review. Considering all 23 manuscripts (including those editorially rejected and not sent for peer review), the mean time from submission to first decision was 4.13 days (Table 1).

**Table 1.** Data on the 23 manuscripts reviewed as part of the Fast & Fair peer review experiment in 2024. All numbers are in days (either total days or working days, where working days excludes weekends and holidays). A value of 0 means less than 24 h. All manuscripts met the goal of reviewed with decisions in 7 working days or less. Outcome column shows first editorial decision. Editorial Reject denotes rejection without peer review.

| Manuscript number | Academic editor before peer review (working days) | Peer review (total days) | Academic editor after peer review (total days) | Total days since submission | Total working days since submission | Outcome |
| --- | --- | --- | --- | --- | --- | --- |
| 1 | 0 | 0 | 0 | 2 | 2 | Editorial Reject |
| 2 | 0 | 6 | 0 | 7 | 5 | Reject |
| 3 | 0 | 6 | 0 | 7 | 5 | Major Revision |
| 4 | 0 | 5 | 0 | 6 | 4 | Reject |
| 5 | 1 | 3 | 0 | 6 | 4 | Reject |
| 6 | 0 | 6 | 0 | 7 | 5 | Reject |
| 7 | 0 | 6 | 0 | 7 | 5 | Major Revision |
| 8 | 0 | 4 | 0 | 5 | 3 | Major Revision |
| 9 | 0 | 1 | 1 | 2 | 2 | Reject |
| 10 | 0 | 0 | 0 | 1 | 1 | Editorial Reject |
| 11 | 0 | 6 | 0 | 7 | 5 | Major Revision |
| 12 | 0 | 6 | 0 | 6 | 4 | Reject |
| 13 | 0 | 6 | 0 | 7 | 5 | Major Revision |
| 14 | 1 | 4 | 0 | 7 | 5 | Minor Revision |
| 15 | 1 | 7 | 0 | 11 | 7 | Major Revision |
| 16 | 0 | 5 | 1 | 7 | 5 | Major Revision |
| 17 | 0 | 3 | 0 | 3 | 3 | Major Revision |
| 18 | 0 | 4 | 0 | 5 | 3 | Major Revision |
| 19 | 0 | 6 | 0 | 8 | 6 | Major Revision |
| 20 | 0 | 6 | 0 | 6 | 4 | Reject |
| 21 | 0 | 6 | 4 | 7 | 5 | Reject |
| 22 | 0 | 4 | 4 | 11 | 7 | Major Revision |
| 23 | 0 | 0 | 0 | 0 | 0 | Editorial Reject |

**Fig 2.**
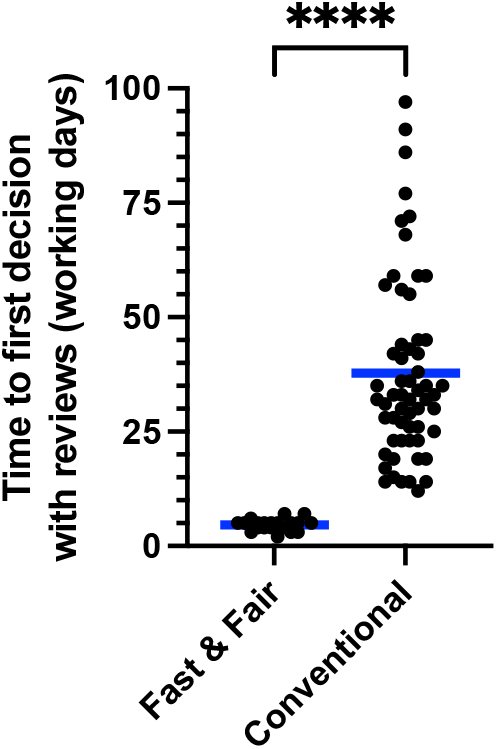
Time to first decisions with reviews for manuscripts submitted July to December, 2024. Each data point is a different manuscript submitted and reviewed as part of the Fast & Fair or conventional peer review process. Manuscripts transferred from other journals with reviews and manuscripts rejected without peer review are not included in this graph. Working days excludes weekends. Blue line is the mean. ****, *p*<0.0001, unpaired *t*-test, two-tailed.

### Editorial assessments of review quality

The review quality was maintained, as measured by how academic editors scored each review. All review reports received a score of 100 (helpful, adequate). No reviews were rated as unhelpful.

Another indicator of editorial assessment and review quality is the rejection rate following first round of peer review. If the rejection rate in the Fast & Fair group was significantly lower than in the control (conventional) review group, this would suggest that Fast & Fair is sacrificing editorial quality for speed. The experimental group (n=20) did not show a statistically significant difference in rejection rate (30%) compared to the control group (n=65, 37%) reviewed in the same time period (chi-square test, χ2=0.321, df=1, *p*=0.76). Therefore, we conclude that in our pilot experiment, rapid review did not occur at the expense of quality of the reviews.

### Retainer versus freelance reviewer models

In total 12 retainer and 68 freelancer reviewers were recruited by the end of the pilot (Table 2). Both sets of reviewers met expectations on review quality and timeliness.

**Table 2.** Reviewers recruited through the Fast & Fair peer review experiment in 2024. Number of recruited reviewers and completed reviews by retainers and freelancer reviewers. Acceptance rate of freelancers shows proportion of invited freelancers that accepted to review.

| Number of retainers recruited | Number of reviews completed by retainers | Number of freelancers recruited | Number of reviews completed by freelancers | Accept rate of freelancers (%) |
| --- | --- | --- | --- | --- |
| 12 | 30 | 68 | 10 | 77 |

The retainer model proved challenging because demand was difficult to estimate, leading to instances where contracted retainers were paid but assigned fewer than three manuscripts per quarter. In addition, this model presented editorial challenges in cases where subject areas were not covered by the small number of retainer reviewers. The freelancer model proved more cost-effective and adaptable to fluctuating manuscript submission rates. As a result of this, at the end of the pilot all retainer reviewers were transitioned to freelancer contracts.

### Costs

In addition to the costs associated with paying reviewers, the Fast & Fair pilot incurred costs associated with setting up the reviewer contracts and administrative costs setting up and maintaining the system to accommodate the quick turnaround of manuscripts (Table 3). We attempted to estimate the additional and unique costs associated with Fast & Fair and exclude existing costs of conventional peer review. The total cost of the 6-month Fast & Fair pilot is estimated to be £50,000-60,000.

**Table 3.**
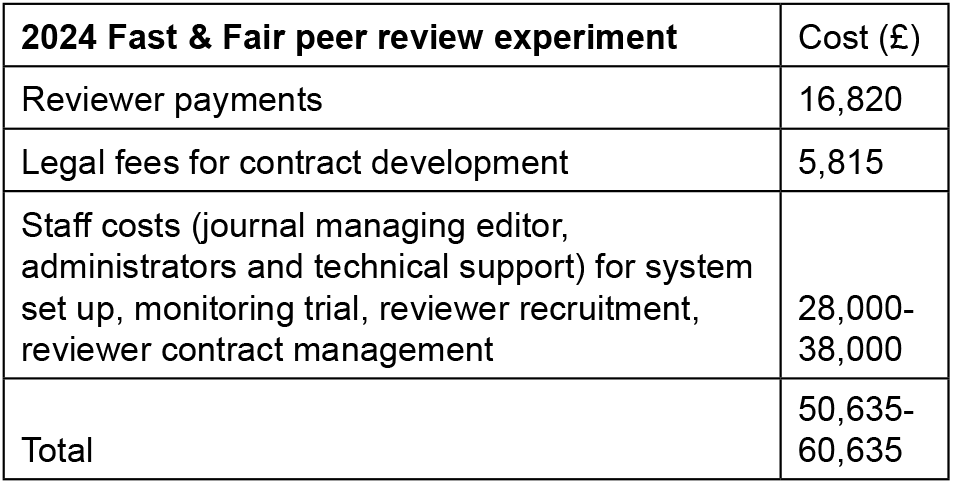
Costs of the Fast & Fair peer review experiment in 2024. For staff costs, we included the upper and lower estimates based on the amount of time staff spent working on Fast & Fair and excluded time spent on other duties, such as supervising conventional peer review.

## DISCUSSION

The Fast & Fair peer review initiative pilot successfully demonstrated that high-quality peer review could be completed within 7 business days. The structured workflow eliminated delays associated with reviewer identification, ensuring that manuscripts were reviewed and returned to authors in a timely manner. This represents a substantial reduction in the average time to first decision compared to the conventional peer review approach (Fig. 2). Maintaining review quality was a priority, and the editorial scoring system confirmed that reviews remained rigorous despite the accelerated timeline. Additionally, the rejection rate following peer review was the same in both Fast & Fair and conventional peer review groups, indicating that the quality of the reviews and editorial assessment was consistent across both approaches. Furthermore, we are not solely reliant on the editorial scoring system; internal oversight is performed where editor editorial staff check reviews to ensure quality.

The recruitment strategy for freelancers was effective and can be adapted to other subjects, which we are currently implementing. To ensure review quality and transparency, reviewer reports will be published alongside accepted manuscripts for papers submitted from January 2025 onward.

One challenge encountered during the Fast & Fair peer review experiment was managing editorial and reviewer absences, particularly during holidays. To mitigate potential delays, we implemented a structured coverage system in which academic editors and reviewers provided advance notice of their availability, allowing us to assign alternative editors or reviewers as needed. This approach ensured that manuscripts continued to progress smoothly and that the 7-business-day turnaround target was consistently met. As we expand Fast & Fair to additional subject areas and a larger editorial team, the complexity of managing coverage will increase. We will continually refine our approach, implementing clear communication and contingency plans to ensure the peer review process remains uncompromised.

How can we reconcile the substantial improvements in review turn-around observed in our Fast & Fair initiative with the more modest effects reported by Cotton et al. (Cotton et al., 2025)? A limitation of Cotton et al. was that financial incentives were provided without explicit performance expectations. Reviewers were offered $250 for completing reviews within the standard timeframe (14 days), without a contract specifying faster turnaround or improved review quality. Cotton et al. attributed their modest improvements (∼1 day faster review) to inherent limits on reviewers’ ability to respond faster in reviewing medical manuscripts. However, several alternative explanations merit consideration. Firstly, absent clear performance expectations, reviewers had little incentive to change their existing practices. By analogy, if an employee receives a raise without clear expectations of enhanced productivity or responsibilities, economic theory predicts no meaningful change in performance – the employee continues performing at the same level, now simply at a higher compensation.

In contrast, our Fast & Fair initiative directly linked financial incentives to explicit reviewer expectations (review completion within 4 business days, adherence to quality standards), which likely motivated reviewers to meet these stricter performance criteria. This explicit linkage of incentives and expectations ensures accountability and could explain why our initiative achieved more substantial improvements in reviewer turnaround compared to the modest effects observed by Cotton et al. Secondly, differences in reviewer demographics and manuscript types may also contribute to the divergent outcomes. Reviewers for Critical Care Medicine may be predominantly physician-scientists, who generally receive higher salaries compared to PhD-level scientists common in biology journals. Consequently, a $250 incentive might represent a relatively smaller motivational factor for physician-scientists compared to PhD-scientist reviewers.

Our experience suggests the Fast & Fair peer review initiative is scalable. Since December 2024, we expanded the reviewer pool to include ∼100 contracted freelancers across various fields. Recruitment efforts were particularly effective by contacting preLighters, which yielded a 70% success rate compared to a 20% success rate from cold emails to scientists recommended by our academic editors. Initially, to select reviewers we relied on the recommendations of our editors to identify scientists with a proven record of producing quality reviews. This pool of reviewers is biased towards scientists who are tenured and who have many years of experience as peer reviewers and/or journal editors. In the future, we aim to have an open call where we identify reviewers based on an application that tests their ability to review a sample manuscript in 4 business days. This would allow us to recruit peer reviewers regardless of career stage, geographic location, or network effect (i.e. who they know).

The Fast & Fair peer review experiment demonstrated that this initiative is feasible for a biology/biomedical journal like Biology Open. As we plan to expand Fast & Fair peer review to other areas of the journal and test its scalability in 2025, we are exploring various models to ensure financial sustainability through 2026.

The pilot’s setup and operational costs were approximately £50,000-60,000, partly due to fixed setup expenses required to establish the new system and the higher costs of the retainer model. However, transitioning retainers to freelancer contracts has helped mitigate these costs. Additionally, fixed expenses, such as legal fees and system setup, are unlikely to recur at scale or in the future. One potential approach to covering reviewer payments is to increase article processing charges (APCs) to authors. AI-assisted workflows may also help reduce publishing costs. We have no plans to increase the APCs for papers published in Biology Open in 2025. In 2026 we will report on the results of the initiative in 2025 and consider possible APC adjustments based on those findings. Unlike large commercial organisations, non-profit publishers like The Company of Biologists neither benefit from economies of scale nor anticipate profits that would cover initiatives like Fast & Fair at scale.

### Limitations of the pilot study

Our pilot study included only 20 manuscripts, so we need larger-scale testing to assess feasibility more broadly. We also focused on developmental biology and physiological sciences, so further evaluation is needed to determine if Fast & Fair works across other biological and biomedical fields.

Predicting the required reviewer expertise was a challenge. We excluded two manuscripts due to gaps in our pre-recruited reviewer pool. Expanding this pool should improve coverage.

Recruiting reviewers was another limitation. Some email invitations may have been lost to spam filters, and some reviewers declined due to the short turnaround time or perceived insufficient compensation. This reduced the number of available reviewers and, in some cases, made it harder to match manuscripts with the right expertise. If we cannot increase the percentage of reviewers who accept our invitations, we will need to extend more invitations or reconsider our compensation structure.

### Limitations of the Fast & Fair approach

A key challenge is the cultural shift required for editors to process manuscripts rapidly. Many are used to a slower workflow, and maintaining this faster system may affect editors’ expectations for reward and recognition to match the higher demands. Although this rapid workflow requires faster decision-making, it is balanced by eliminating the time spent finding willing reviewers.

Frequent reviewers may also face fatigue, requiring continuous recruitment to maintain a strong pool. Additionally, the Fast & Fair model introduces the risk of motivation crowding, where financial incentives could reduce intrinsic motivation over time, potentially affecting reviewer retention.

Another concern is the potential increase in low-quality submissions, particularly from paper mills, attracted by faster processing times. We need safeguards to filter out unsuitable manuscripts early while ensuring peer review remains rigorous despite the compressed timeline.

## Conclusion

The Fast & Fair peer review pilot experiment successfully demonstrated that rapid, high-quality peer review is feasible in a biology/ biomedical journal. Eliminating the reviewer identification bottleneck significantly improved efficiency for academic editors, and the freelancer model proved to be a more cost-effective solution for matching reviewers to manuscripts based on expertise compared to the retainer model. Future efforts will focus on expanding the Fast & Fair peer review initiative across additional subject areas, refining editorial workflows, and exploring the financial sustainability of the initiative at scale. By offering an efficient alternative to conventional peer review, the Fast & Fair peer review initiative represents a transformative model for academic publishing.

## Acknowledgments

We thank our academic editors, reviewers, and The Company of Biologists staff including Sue Chamberlain, Laura Patterson, Ania Crowther, Laura Tolhurst, Laura Mason, Andrea Kendall, Lindsey Cole, and Rachel Hackett for their invaluable assistance. We thank Reinier Prosee for helping to recruit preLighters as reviewers. We also thank the Board of Directors at The Company of Biologists for their support, especially the members assigned to advise Biology Open: Jane Langdale, Steven Royle, Laura Machesky and Daniel St Johnston. We are grateful to Claire Moulton for advice throughout the experiment and Katherine Brown and Claire Moulton for comments on the manuscript.

## Competing interests

The authors of this manuscript have the following competing interests: D.A.G. is the Editor-in-Chief of Biology Open and A.C. is the Managing Editor of Biology Open.

## References

Chetty, R., Saez, E. and Sándor, L. (2014). What Policies Increase Prosocial Behavior? An Experiment with Referees at the Journal of Public Economics. Journal of Economic Perspectives 28, 169–188.

Cotton, C. S., Alam, A., Tosta, S., Buchman, T. G. and Maslove, D. M. (2025). Effect of Monetary Incentives on Peer Review Acceptance and Completion: A Quasi-Randomized Interventional Trial. Crit. Care Med.

Doble, E., Schroter, S., Price, A. and Abbasi, K. (2024). The BMJ will remunerate patient and public reviewers. BMJ 387, q2581.

Frey, B. S. and Jegen, R. (2001). Motivation Crowding Theory. J. Econ. Surv. 15, 589–611.

Nguyen, V. M., Haddaway, N. R., Gutowsky, L. F. G., Wilson, A. D. M., Gallagher, A. J., Donaldson, M. R., Hammerschlag, N. and Cooke, S. J. (2015). How long is too long in contemporary peer review? Perspectives from authors publishing in conservation biology journals. PLoS ONE 10, e0132557.

Preston, A. and Johnston, D. (2013). The Future of Academic Research. Figshare.

Ravindran, S. (2016). Getting credit for peer review. Science.

Yu, H., Liang, Y. and Xie, Y. (2024). Can peer review accolade awards motivate reviewers? A large-scale quasi-natural experiment. Humanit. Soc. Sci. Commun. 11, 1557.

